# Homologous expression and characterization of gassericin T and gassericin S, a novel class IIb bacteriocin produced by *Lactobacillus gasseri* LA327

**DOI:** 10.1101/435081

**Authors:** Genki Kasuga, Masaru Tanaka, Yuki Harada, Hiroshi Nagashima, Taisei Yamato, Ayaka Wakimoto, Yoshiyuki Ito, Yasushi Kawai, Jan Kok, Testuya Masuda

**Author notes:** Address correspondence to Yasushi Kawai.

## Abstract

*Lactobacillus gasseri* LA327 isolated from the large intestine tissue in humans is a bacteriocinogenic strain and is predicted to produce two kinds of class IIb bacteriocins, i.e. gassericin T (GT) and acidocin LF221A (Acd LF221A). In this study, DNA sequencing of the genes for GT and Acd LF221A on *Lb. gasseri* LA327 revealed that the amino acid sequences for GT completely corresponded with those of *gat* except for GatK (histidine kinase). However, those for the Acd LF221A had analogues which differed in at least one amino acid residue to be a putative class IIb bacteriocin designated as gassericin S (GS). By deletion test of GT structural genes (*gatAX*), the LA327 strain retained the bacteriocin activity, and the LA327 mutant strain lacking the ABC-type transporter gene (*gatT*) completely lost the bacteriocin activity. This indicates that LA327 strain is a GS producer, and GS production is performed via *gat* with the inclusion of *gatT*. Homologous expression using deletion mutants for GS and GT containing each single peptide elucidated that GS (GasAX) and GT (GatAX) showed synergistic activity as class IIb bacteriocins, respectively, and no synergistic activity was observed between each peptide of GS and GT. The molecular mass of GS was estimated to be theoretical ca. 5,400 Da by *in situ* activity assay after SDS-PAGE, clarifying that GS was actually expressed as an active class IIb bacteriocin. Furthermore, stability of GS expressed against pH, heat and protease was determined.

**Importance:** We determined the complete DNA sequence for GS, a novel class IIb bacteriocin of *Lb. gasseri*, and succeeded to express GS as active bacteriocins. Our results clarified the interaction of each class IIb component peptide for GT in addition to GS via construction of homologous mutants which were not dependent on the purification. These data may demonstrate the characteristics of class IIb bacteriocins for *Lb. gasseri*.

## Introduction

Lactic acid bacteria (LAB), one of the potent candidates which are generally recognized as safe (GRAS) microorganisms, have a long history of consumption by humans, and inhabiting various ecosystems, including gastrointestinal (GI) tracts of humans and animals, and other environments (for instance, vegetables, milks, and meats). In addition, lactobacilli are recognized as important members of beneficial GI microbiota of humans and animals, along with bifidobacteria ^1^. Especially, *Lactobacillus gasseri* is one of the predominant species in lactobacilli of human small intestines, and has been isolated not only from the GI tracts and the feces, but also the oral and the vaginal cavities and mammary areola, and is considered one of the representative probiotics ^2–5^.

Many researchers have reported the beneficial effects of *Lb. gasseri* on the hosts, including the immunoregulation, the alleviation of allergic symptoms, the prevention of bacterial and viral infections, the antitumor effect, and the inhibition of lipid absorption ^6–11^. Indeed, many yogurt products containing *Lb. gasseri* strains are commercially sold in Japan to provide the benefits mentioned above to consumers. It has been known that LAB may produce various antimicrobial agents, such as organic acids, diacetyl, hydrogen peroxide, and bacteriocins, to survive in competitive microbial niches.

Bacteriocin is defined as ribosomal synthesized antimicrobial peptides and proteins from microorganisms, and it is expected as a potential alternative of antibiotics due to the several advantageous properties, including their potency (as determined *in vitro* and *in vivo*), variety of antimicrobial spectra (both narrow and broad), low toxicity, possibility of *in situ* production by probiotics, and the fact that these peptides can be bioengineered ^12–13^. Presently, bacteriocins from Gram-positive bacteria are classified into two major classes of lantibiotics (class I, bacteriocins containing post-translationally modified residues) and non lantibiotics (class II, bacteriocins with non-modified residues, excluding the formation of disulfide bridges and circular bacteriocins), and class II is subclassified into four subclasses of class IIa (pediocin-like), IIb (two-peptide), IIc (cyclic), and IId (not correspond to IIa-IIc) ^13–14^.

Previously, we reported that gassericin A (GA), a head-to-tail cyclized bacteriocin (class IIc) from *Lb. gasseri* LA39 ^15^ and gassericin T (GT), a putative two-peptide bacteriocin (class IIb) consisted of GatA and GatX from *Lb. gasseri* SBT2055 and LA158 ^16–17^. In addition, Bogovic-Matijasic *et al.* (1998) ^18^ reported that acidocin LF221A (Acd LF221A) and acidocin LF221B (Acd LF221B) were produced by *Lb. gasseri* LF221: the latter was analogue of GatX (the second peptide of GT) containing alanine at position of 50 instead of glycine in GatX, and the former was potentially a novel bacteriocin derived from *Lb. gasseri*. Recently, we isolated *Lb. gasseri* LA327, a producer strain of two bacteriocins estimated as GT and Acd LF221A, from the human large intestine tissue. As other analogues of GT, gassericin K7B from *Lb. gasseri* K7 and gassericin E from *Lb. gasseri* EV1461, and as completely homolog of Acd LF221A, gassericin K7A from *Lb. gasseri* K7, have been reported as class IIb bacteriocins of *Lb. gasseri* ^19–20^. However, there have never been any reports for success of complete purification of class IIb bacteriocins produced by *Lb. gasseri*, and the synergistic activity by coexistence of each peptide, which should be an essential character for class IIb bacteriocin, is not still demonstrated.

In this study, we focused on identification and characterization of two class IIb bacteriocins detected from *Lb. gasseri* LA327 and demonstration of the two-component action through homologous expression of these bacteriocins using genetic constructed mutant strains.

## Results

### DNA sequencing and genetic analysis

The nucleotide sequencing surrounding *gatAX* and *gasAX* (like acidocin LF221A and gassericin K7A) encoded in chromosomal DNA of *Lb. gasseri* LA327 decided 6,935 bp and 1,143 bp DNA sequences harboring nine and three ORFs (DDBJ accession No. LC389592 and LC389591), respectively. The homology searching of predicted amino acid sequences for peptides and proteins encoded in each ORFs revealed that the nine products for *gat* accorded to those of *Lb. gasseri* LA158 (GenBank Accession No. AB710328) except one amino acid residue in GatK (amino acid residue at the position 168 is valine in LA327 instead of alanine in LA158, data not shown), and the three ORFs for *gas* were not detected in LA158. Furthermore, *gat* and *gas* were similar to that of acidocin LF221B (Acd LF221B), gassericin K7B, and that of acidocin LF221A (Acd LF221A) and gassericin K7A, respectively (Table 1-2). However, the sequences of two class IIb bacteriocins for *gas* and *gat* in *Lb. gasseri* LA327 were slightly different from Acd LF221A and Acd LF221B in the range of 0-2 amino acid residues (GenBank Accession No. AY295874.1 and AY297947.1) (Fig. 1ab), naming the new putative bacteriocin as gassericin S (GasA and GasX).

**Table 1.** Deduced peptides and proteins encoded in *gat* from *Lactobacillus gasseri* LA327.

| ORF | Length<br>(amino acid) | MW | pI | Function | Homology search<sup>a</sup><br>(identities: residual ratio, %) | Localization<br>(TMS no.)<sup>b</sup> |
| --- | --- | --- | --- | --- | --- | --- |
| *gatP* | 50 | 5,852 | 9.83 | inducer peptide precursor | lactacin F inducer peptide precursor<br>*Lb. gasseri* K7 (50/50,100%) | soluble<br>(none) |
| *gatK* | 435 | 50,444 | 5.2 | histidine kinase | histidine kinase *Lb. gasseri* K7<br>(430/435,99%) | membrane<br>(6) |
| *gatR* | 265 | 30,280 | 5.36 | response regulator | chemotaxis protein CheY<br>*Lb. gasseri* K7 (265/265,100%) | soluble<br>(none) |
| *gatT* | 719 | 81,054 | 7.71 | ABC transporter permease component | peptide ABC transporter ATP-binding protein<br>*Lb. gasseri* K7 (718/719,99%) | membrane<br>(6) |
| *gatC* | 197 | 21,938 | 10.13 | putative accessory protein | putative gassericin K7 B accessory protein<br>(196/197,99%) | membrane<br>(1) |
| *gatZ* | 33 | 3,924 | 7.57 | putative uncharacterized protein | unknown *Lb. gasseri* LF221<br>(33/33,100%) | soluble<br>(none) |
| *gatA* | 75 | 7,480 | 10.02 | GatA precursor | putative complement factor Acd221b<br>*Lb. gasseri* LF221 (75/75,100%) | membrane<br>(1) |
| *gatX* | 65 | 6,673 | 10.11 | GatX precursor | acidocin LF221B Acd221B<br>*Lb. gasseri* LF221 (64/65,98%) | soluble<br>(none) |
| *gatI* | 112 | 13,343 | 9.81 | immunity protein | putative immunity protein Aci221B<br>*Lb. gasseri* LF221 (112/112,100%) | membrane<br>(4) |

<sup>a</sup> Homology search was demonstrated with BLAST program
(http://blast.ddbj.nig.ac.jp/top-j.html).
<sup>b</sup> Deduced using the TMHMM program (http://www.cbs.dtu.dk/services/TMHMM/).

**Table 2.** Deduced peptides and proteins encoded in *gas* from *Lactobacillus gasseri* LA327.

| ORF | Length<br>(amino acid) | MW | pI | Function | Homology search<sup>a</sup><br>(identities: residual ratio, %) | Localization<br>(TMS no.)<sup>b</sup> |
| --- | --- | --- | --- | --- | --- | --- |
| *gasA* | 79 | 7,662 | 10.27 | GasA precursor | gassericin K7 A complemental<br>factor (78/79,99%) | membrane<br>(1)<sup>c</sup> |
| *gasX* | 69 | 7,287 | 10.00 | GasX precursor | acidocin LF221A Acd221A<br>(67/69,97%) | membrane<br>(1)<sup>c</sup> |
| *gasI* | 90 | 10,789 | 7.42 | putative<br>immunity protein | gassericin K7 A immunity<br>protein (90/90,100%) | membrane<br>(2) |

<sup>a</sup> Homology search was demonstrated with BLAST program
(http://blast.ddbj.nig.ac.jp/top-j.html).
<sup>b</sup> Deduced using the TMHMM program (http://www.cbs.dtu.dk/services/TMHMM/).
<sup>c</sup> With putative mature peptide.

**Fig. 1a.**
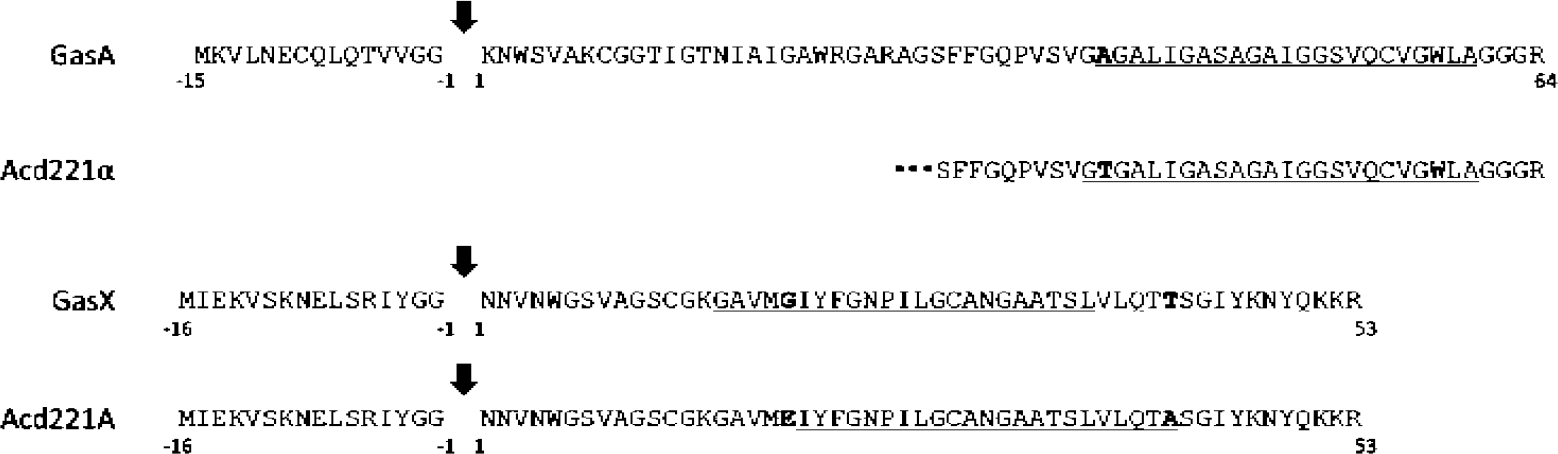
Comparison of deduced amino acid sequences for gassericin S (GasA and GasX) and acidocin LF221A (Acd221α and Acd221A). Arrows, underlines, and bolds indicate predicted cleavage sites, putative transmembrane domains, and different amino acid residues between GS and Acd LF221A, respectively.

**Fig. 1b.**
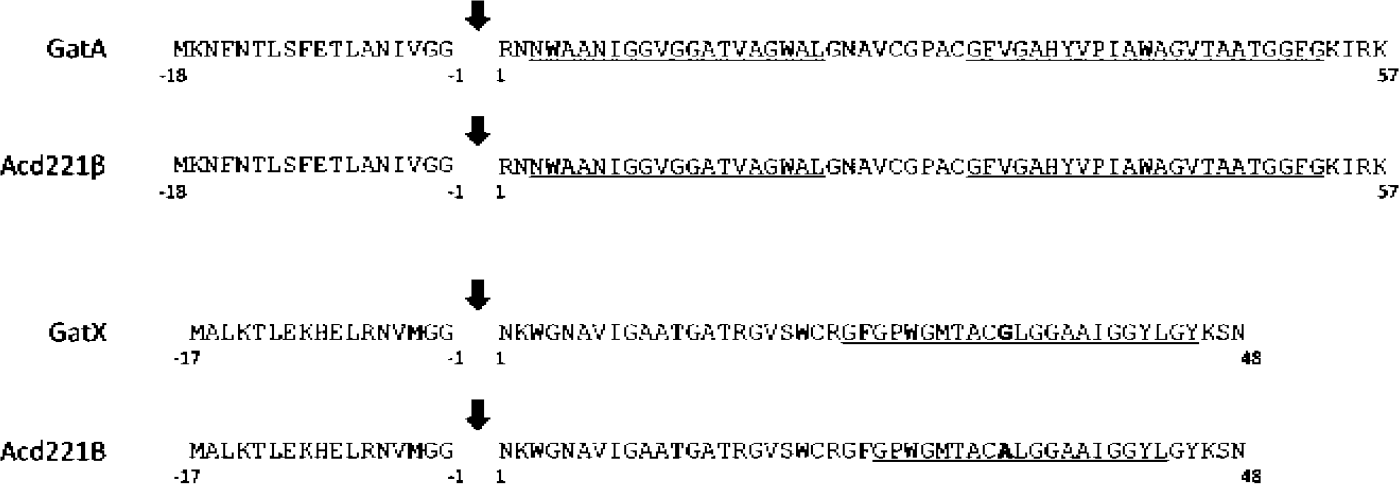
Comparison of deduced amino acid sequences for gassericin T (GatA and GatX) and acidocin LF221B (Acd221β and Acd221B). Arrows, underlines, and bolds indicate predicted cleavage sites, putative transmembrane domains, and different amino acid residues between GT and Acd LF221B, respectively.

### Verification of gassericin S production mechanism

GT activity (15,754 AU/mL) in the culture supernatant of *Lb. gasseri* LA158 was completely lost by deletion of the GT structural gene (*gatAX*) from chromosomal DNA of LA158 (LA158⊿*gatAX*), and *Lb. gasseri* LA327 kept and raised bacteriocin activity (from 16 AU/mL to 64 AU/mL) after the *gatAX* was eliminated (LA327⊿*gatAX*). Continuously, the bacteriocin activity in the culture supernatant of LA327 vanished by elimination of the putative ABC-transporter (GatT) from chromosomal DNA of LA327 (LA327⊿*gatT*), suggesting that GS is a bacteriocin secreted thorough GatT.

### Synergistic activity among each peptide of gassericin S and gassericin T

Bacteriocin activity (62 AU/mL and 123 AU/mL) was detected in the culture supernatants of co-producer strains for GasAX, *Lb. gasseri* LA158⊿*gatAX* (pGS-AX) and (pGS-AXI), respectively, and no activities were obtained in the culture supernatants of single-producer strains for GasA and GasX, *Lb. gasseri* LA158 ⊿ *gatAX* (pGS-AX ⊿X), (pGS-AXI⊿X), (pGS-AX⊿A), and (pGS-AXI⊿A), even in concentrated forty-folds (Table 3). On the other hand, mixing GasA (GasAI) and GasX (GasXI) with equal amount showed synergistic activity (Table 3).

**Table 3.**
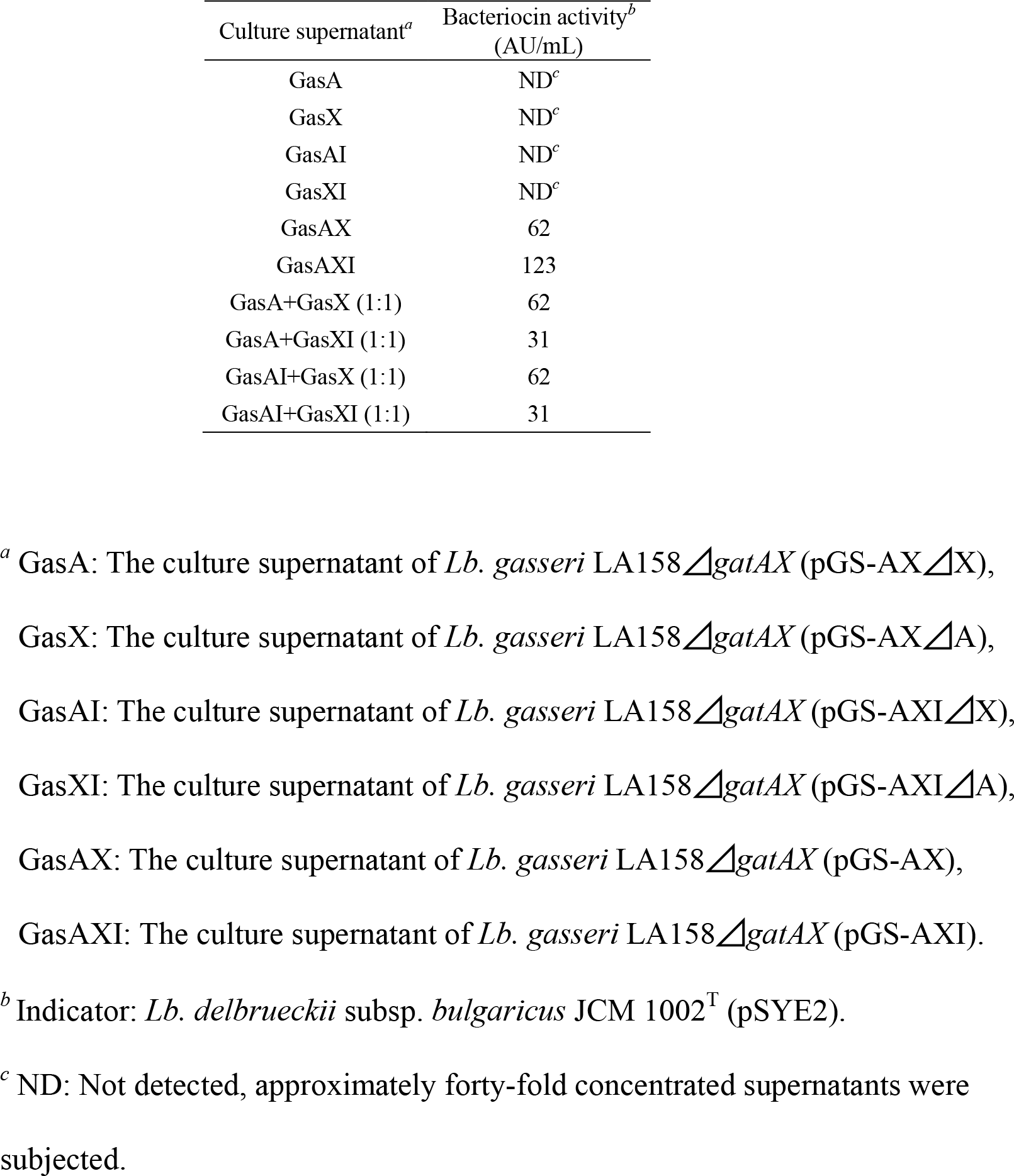
Antibacterial activities of each Gas peptide in alone and combination.

| Culture supernatant<sup>a</sup> | Bacteriocin activity<sup>b</sup><br>(AU/mL) |
| --- | --- |
| GasA | ND<sup>c</sup> |
| GasX | ND<sup>c</sup> |
| GasAI | ND<sup>c</sup> |
| GasXI | ND<sup>c</sup> |
| GasAX | 62 |
| GasAXI | 123 |
| GasA+GasX (1:1) | 62 |
| GasA+GasXI (1:1) | 31 |
| GasAI+GasX (1:1) | 62 |
| GasAI+GasXI (1:1) | 31 |

<sup>a</sup> GasA: The culture supernatant of *Lb. gasseri* LA158 $\Delta$ gatAX (pGS-AX $\Delta$ X), GasX: The culture supernatant of *Lb. gasseri* LA158 $\Delta$ gatAX (pGS-AX $\Delta$ A), GasAI: The culture supernatant of *Lb. gasseri* LA158 $\Delta$ gatAX (pGS-AXI $\Delta$ X), GasXI: The culture supernatant of *Lb. gasseri* LA158 $\Delta$ gatAX (pGS-AXI $\Delta$ A), GasAX: The culture supernatant of *Lb. gasseri* LA158 $\Delta$ gatAX (pGS-AX), GasAXI: The culture supernatant of *Lb. gasseri* LA158 $\Delta$ gatAX (pGS-AXI).
<sup>b</sup> Indicator: *Lb. delbrueckii* subsp. *bulgaricus* JCM 1002<sup>T</sup> (pSYE2).
<sup>c</sup> ND: Not detected, approximately forty-fold concentrated supernatants were subjected.

Similarly, the assay for synergistic activity of component peptides of GT (GatA and GatX) in the culture supernatants of single-producer strains for GatA and GatX, *Lb. gasseri* LA158⊿*gatX* and ⊿*gatA*, clarified that GatA displayed activity (62 AU/mL) when alone, and remarkably high activity (1,969 AU/mL) was observed by combination of GatA and GatX (Table 4). In the synergistic activity assay between each GS and GT component peptide, except for the original combination of GS and GT (GasA-GasX, and GatA-GatX), no synergistic activity was observed (Table 5).

**Table 4.**
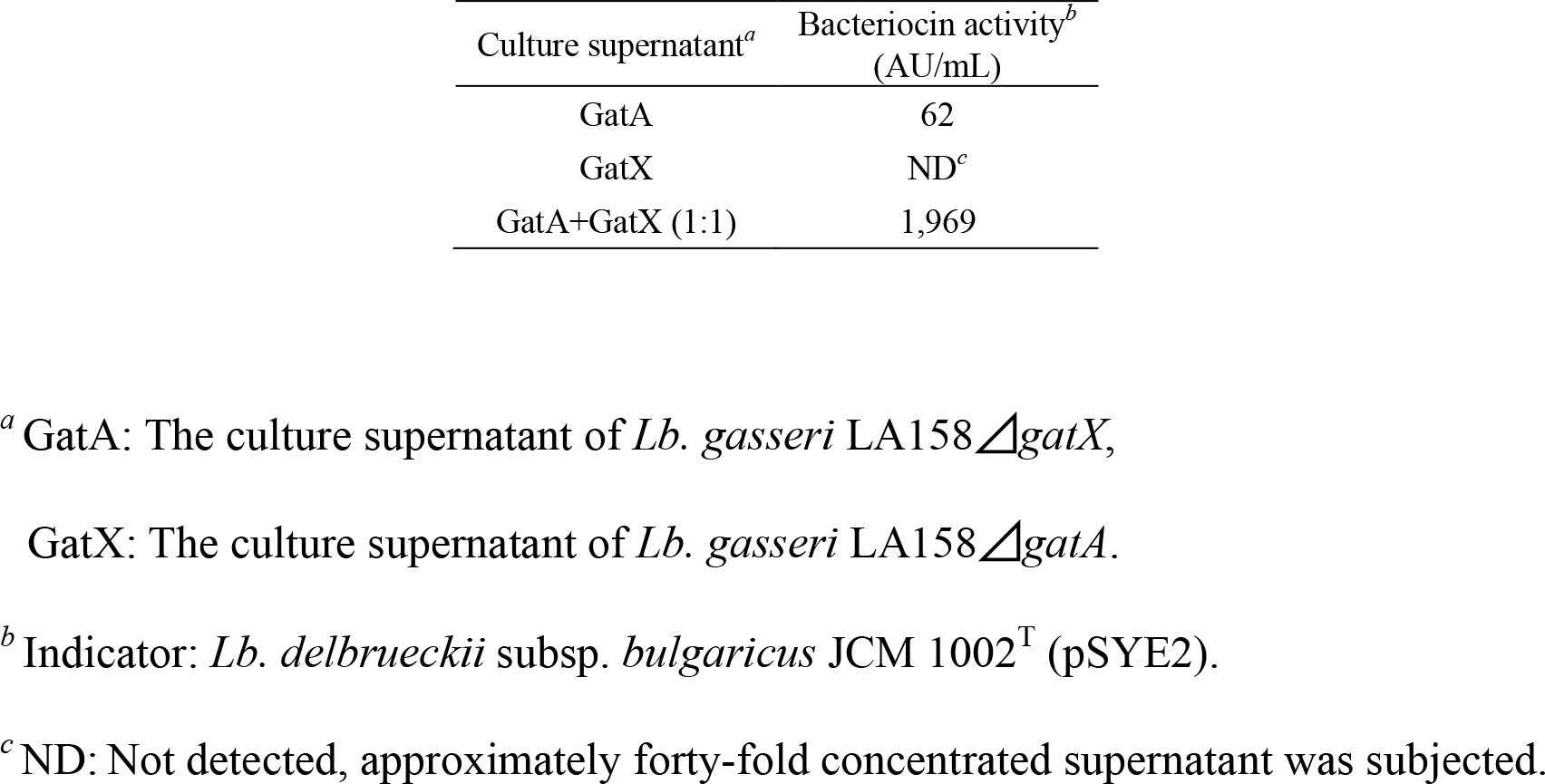
Synergistic activity of GatA and GatX

**Table 5.**
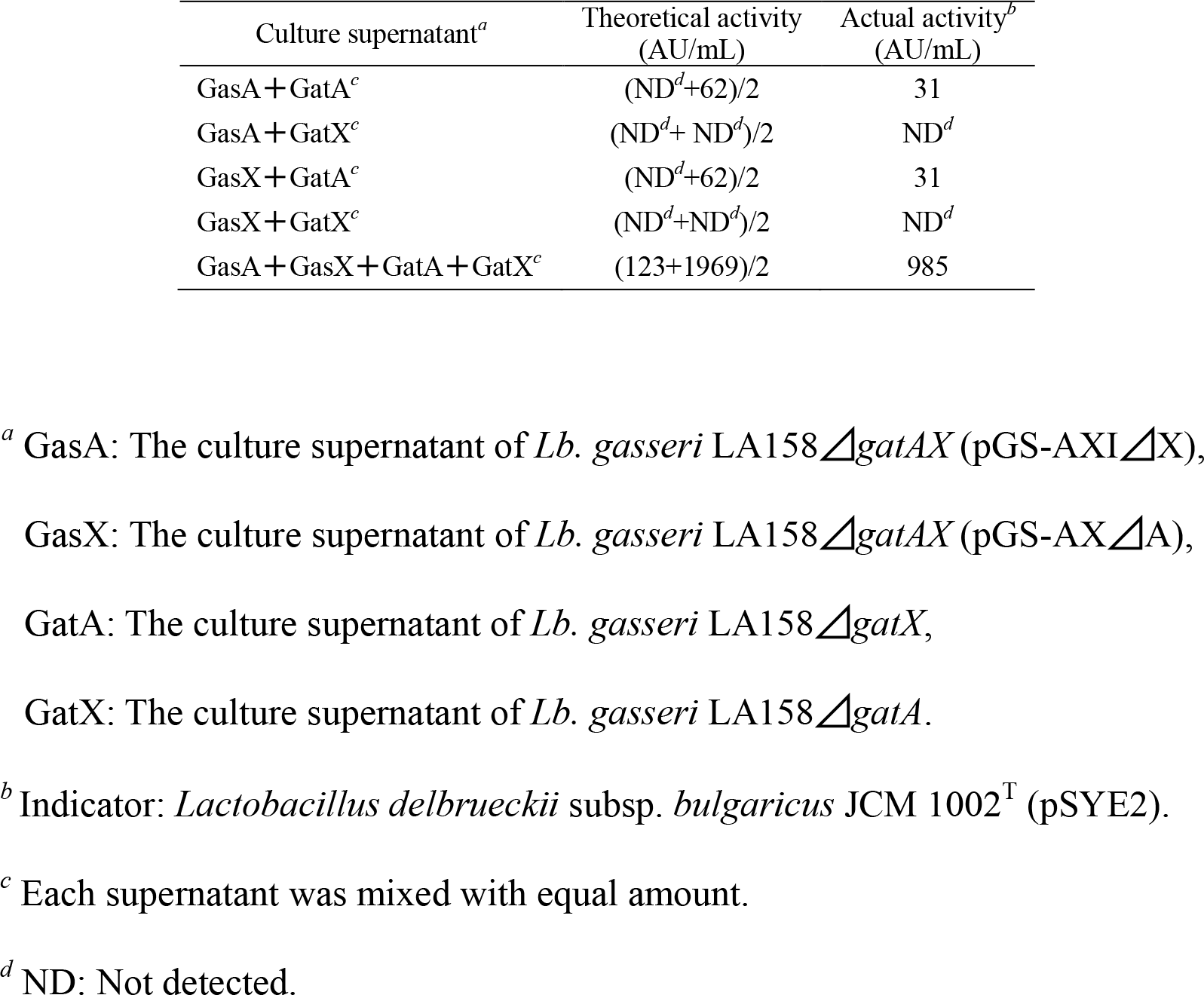
Synergistic activity assay of GS and GT components.

| Culture supernatant<sup>a</sup> | Theoretical activity<br>(AU/mL) | Actual activity<sup>b</sup><br>(AU/mL) |
| --- | --- | --- |
| GasA + GatA<sup>c</sup> | (ND<sup>d</sup>+62)/2 | 31 |
| GasA + GatX<sup>c</sup> | (ND<sup>d</sup>+ ND<sup>d</sup>)/2 | ND<sup>d</sup> |
| GasX + GatA<sup>c</sup> | (ND<sup>d</sup>+62)/2 | 31 |
| GasX + GatX<sup>c</sup> | (ND<sup>d</sup>+ND<sup>d</sup>)/2 | ND<sup>d</sup> |
| GasA + GasX + GatA + GatX<sup>c</sup> | (123+1969)/2 | 985 |

<sup>a</sup> GasA: The culture supernatant of *Lb. gasseri* LA158 $\Delta$ gatAX (pGS-AXI $\Delta$ X),
GasX: The culture supernatant of *Lb. gasseri* LA158 $\Delta$ gatAX (pGS-AX $\Delta$ A),
GatA: The culture supernatant of *Lb. gasseri* LA158 $\Delta$ gatX,
GatX: The culture supernatant of *Lb. gasseri* LA158 $\Delta$ gatA.
<sup>c</sup> Each supernatant was mixed with equal amount.
<sup>d</sup> ND: Not detected.

### *In situ* activity assay for gassericin S

A clear zone derived with GS appeared by the *in situ* activity assay after SDS-PAGE and the molecular mass of GS was presumed as approx. 5,400 Da by standard curve constructed with electrophoresis distance of each molecular marker (Fig. 2).

**Fig. 2.**
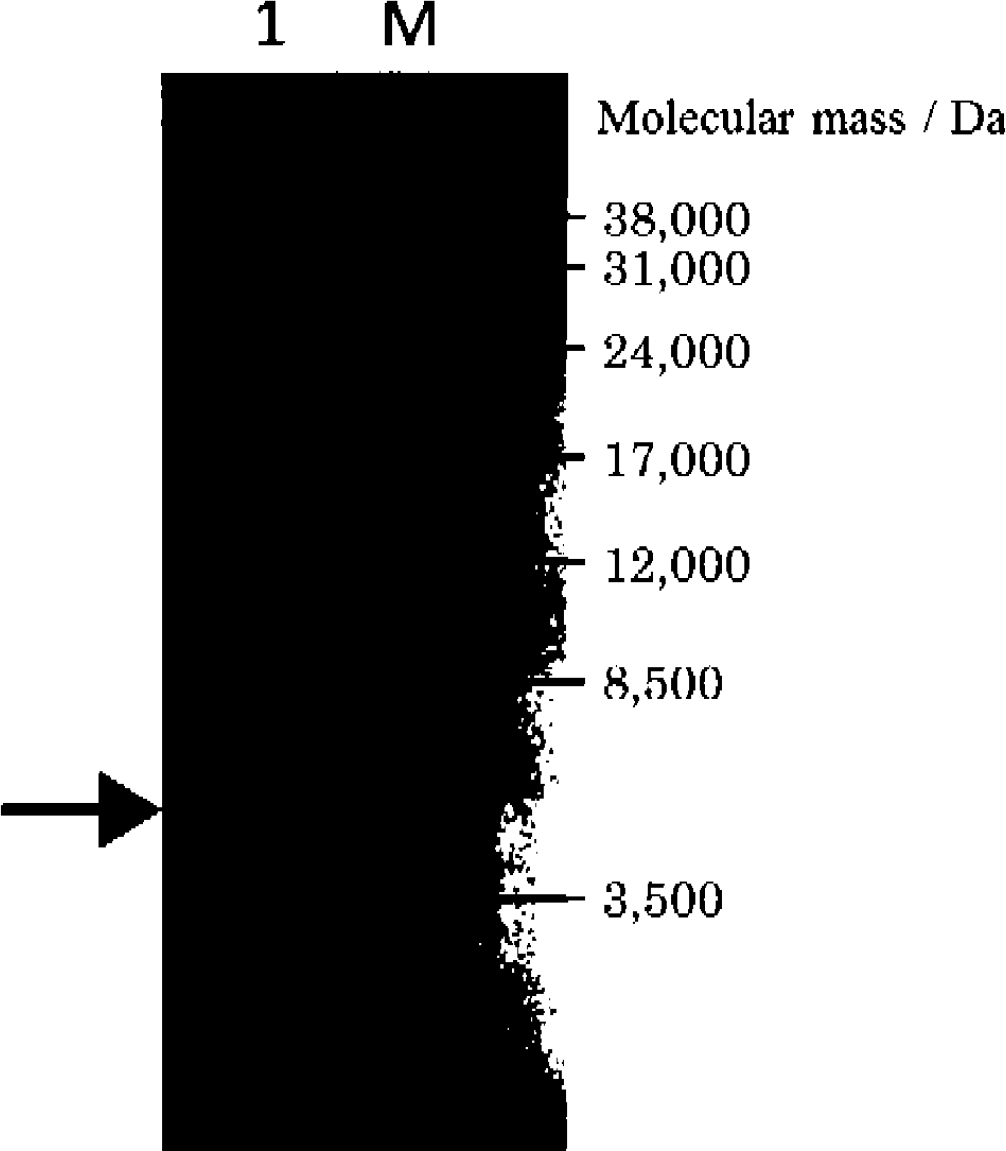
*In situ* activity assay of semi-purified gassericin S. Indicator: *Lactobacillus delbrueckii* subsp. *bulgaricus* JCM 1002^T^ (pSYE2). Lane M, molecular weight markers; lane1, semi-purified gassericin S. Allow indicates the clear zone of gassericin S.

### pH, heat, and proteolysis stability of gassericin S

GS stability under various pH, thermal, and proteolytic conditions was investigated. The bactetiocin activity of GS in the culture supernatant was stable in wide pH range (pH 2, 4, 7, 10) (data not shown) and 25% of GS activity was retained in the most severe heating condition (121 °C, 15 min) in the tests (Table 6). The GS conservation test elucidates that the GS activity decreased to 25% by incubation at 37 °C, but was entirely maintained at 4 °C (Table 6). By examination of proteolytic resistance for GA, GT, and GS, there were no bacteriocins showing tolerance against proteinase K. GS completely lost the activity by all proteases used in this study, and GT had slight tolerance only for pepsin. On the other hand, the bacterioicin activity of GA, a circular bactericidal peptide, was detected after treatment of all proteases used, except for proteinase K (Table 7).

**Table 6.** Heat stability of gassericin S^*a*^.

| Condition | Bacteriocin activity<sup>b</sup><br>(AU/mL, residual rate) |
| --- | --- |
| untreated | 123 |
| 121°C, 15 min | 31 (25%) |
| 95°C, 5 min | 62 (50%) |
| 70°C, 1 h | 62 (50%) |
| 37°C, 1 mth | 31 (25%) |
| 4°C, 1 mth | 123 (100%) |

<sup>a</sup> The culture supernatant of *Lb. gasseri* LA158 $\Delta$ gatAX (pGS-AXI).
<sup>b</sup> Indicator: *Lactobacillus delbrueckii* subsp. *bulgaricus* JCM 1002<sup>T</sup> (pSYE2).

**Table 7.** Proteolytic resistant of bacteriocins produced by *Lb. gasseri* tested.

| Protease | Residual activity of bacteriocins (AU/mL, rate)<sup>a</sup> |  |  |
| --- | --- | --- | --- |
|  | GA<sup>b</sup> | GT<sup>c</sup> | GS<sup>d</sup> |
| untreated | 985 | 31,508 | 246 |
| pepsin | 492 (50%) | 492 (1.6%) | ND<sup>e</sup> |
| trypsin | 62 (6.3%) | ND<sup>e</sup> | ND<sup>e</sup> |
| alpha-chymotrypsin | 246 (25%) | ND<sup>e</sup> | ND<sup>e</sup> |
| proteinase K | ND<sup>e</sup> | ND<sup>e</sup> | ND<sup>e</sup> |

<sup>a</sup> Indicator: *Lb. delbrueckii* subsp. *bulgaricus* JCM 1002<sup>T</sup> (pSYE2).
<sup>b</sup> GA: gassericin A, the culture supernatant of *Lb. gasseri* LA39
<sup>c</sup> GT: gassericin T, the culture supernatant of *Lb. gasseri* LA158
<sup>d</sup> GS: gassericin S, the culture supernatant of *Lb. gasseri* LA327
<sup>e</sup> ND: Not detected.

## Discussion

DNA sequencing of the genes for GT and Acd LF221A on *Lb. gasseri* LA327 revealed 6,935 bp and 1,143 bp DNA sequences containing 9 ORFs and 3 ORFs (DDBJ accession No. LC389592 and LC389591), respectively. The amino acid sequences derived from 9 ORFs for GT were completely in accordance with those of *gat* (from LA158, GenBank Accession No. AB710328) except for GatK (histidine kinase, A168V) and *Lb. gasseri* LA327 may be a GT producer. The amino acid sequences of three ORFs for Acd LF221A were the analogue differing in at least one amino acid residue (Fig. 1a) and the putative class IIb bacteriocin which was designated as gassericin S (GS).

The primer walking analysis elucidated that no genes related to regulation and transporting for GS, a two-component bacteriocin, were recognized in the surrounding areas of *gas* (the gene cluster of GS) (data not shown), suggesting that the GS cluster was constructed with only three genes, *gasAX* (structural genes) and *gasI* (immunity gene) and GS may be produced using a secretion system on the LA327 genome.

By deletion test of GT structural genes (*gatAX*) from the *Lb. gasseri* LA158 (GT producer) and *Lb. gasseri* LA327 (putative GT/GS producer), LA327 strain only retained the bacteriocin activity unlike LA158 strain. The LA327 mutant strain lacking the ABC-type transporter gene (*gatT*) completely lost all bacteriocin activity. These results indicate that LA327 strain is a GS producer and GS production is performed via *gat* including *gatT*.

Class IIb bacteriocins exert synergistic activity of each peptide that has no and/or weak activity. Complete purification of both peptides for class IIb bacteriocins from *Lb. gasseri* has never been succeed, and the certification of synergistic activity mentioned above is still unclarified in gassericins. In this study, we firstly demonstrated that GS and GT consisting of GasAX and GatAX, respectively, showed synergistic activity as class IIb bacteriocins by homologous expression containing deletion mutants of each single peptide. It was also elucidated that GatA has antimicrobial activity when alone differing from the other peptides (GatX, GasA, and GasX, even if concentrated forty-fold), and no synergistic activity was observed between each peptide of GasAX and GatAX against the indicator strain, *Lb. delbreuckii* subsp. *bulgaricus* JCM 1002^T^ (Table 3-5). The molecular mass (ca. 5,400 Da) of GS estimated by *in situ* activity assay after SDS-PAGE was similar to those of GasA and GasX (6,061 Da and 5,481 Da, respectively), clarifying that GS was actually expressed as an active class IIb bacteriocin.

Majhenic *et al.* (2004) ^21^ reported that Acd221A and Acd221B isolated from *Lb. gasseri* LF221, two or one amino acid variants of GasX and GatX, respectively, inhibited the growth of *Lb. sakei* NCDO2714 when alone. Although the difference of the indicator strains used may be related to the results, it was more likely due to the variation of the amino acid residues on Acd221A and Acd221B. Influences of single amino acid variation between second peptides of enterocin C and enterocin 1017 against to their antimicrobial spectra were clarified ^22^. However, the inhibition ability against *Lb. delbrueckii* subsp. *bulgaricus* LMG 6901^T^ (= JCM 1002^T^) of Gas K7_Acp, 100% homolog of GatX produced by *Lb. gasseri* K7, was reported ^23^, indicating that the structure of Gas K7_Acp may be different from inactive GatX in this study, owing to test solutions and/or expression hosts used, in spite of the same indicator strain and amino acid sequences. Each component peptide (CbnX and CbnY) of class IIb bacteriocin carnobacteriocin XY presented high α-helix content in trifluoroethanol solution, while CbnXY did not show remarkable second structures in aqueous conditions ^24^. Furthermore, our previous study for isolation and purification of GatA and GatX from the culture supernatant of *Lb. gasseri* LA158 failed because of polymerization of GatA and GatX in 60% 2-propanol at the final step for HPLC purification ^17^. Therefore, the organic solvents used for bacteriocin extraction may be involved in activity exhibition of bacteriocins such as GatX and their analogues reported. In contrast, an active peptide of GaeE (one amino acid variants of GatA, W22L), a component peptide of gassericin E produced by *Lb. gasseri* EV1461 ^20^, supports that GatA has the antibacterial activity in alone. Moreover, in this study, a putative immunity protein, GasI, would promote the production of GasAX, while each GasA and GasX production was not affected by *gasI* expression (Table 3). Ra *et al.* (1991) ^25^ reported that deletion of *nisI*, the immunity protein of nisin A, led decrease of nisin production. These results and information indicate that resistant ability of GasI may be unnecessary for production of each inactive single peptide for GasAX, while GasI supports the active GasAX production.

It is not rare that multiple bacteriocins are produced by single strain, but many questions still remain about its advantage. In this study, the synergistic activity was obtained between each complemental peptide of GT and GS and not shown in cross-combination of GT and GS (Table 5). Similarly, no synergistic activity was observed in cross-combination of plantaricin EF and plantaricin JK ^26^, but high activity was obtained in the case of hemilateral peptide of lactococcin G with the complemental peptide of enterocin 1017 and lactococcin Q ^27–29^. It seems that presentation of cross-combination activity is dependent on amino acid sequence similarity for lowly GS-GT (ca. 26%, calculated by BLAST; https://blast.ncbi.nlm.nih.gov./Blast.cgi) and high lactococcin G and enterocin 1017/lactococcin Q (ca. 88% and 57%, respectively).

After treatment tests at various pH and temperatures, the bacteriocin activity of GS produced by *Lb. gasseri* LA158⊿*gatAX* (pGS-AXI) remained, even when exposed in pH 2-10 and heated to 121 °C for 15 min (Table 6-7). However, the activity was remarkably decreased by the incubation at 37 °C for long term, differing from complete maintenance at 4 °C (Table 6). In our previous study, the bacteriocin activity of GT produced by LA158 similarly decreased during 37 °C incubation, but the decay rate of the activity was weakened after heat treatment of the GT solution at 99 °C for 10 min (data not shown), suggesting that the degradation of GT and GS activity may be caused by extracellular protease(s) derived from LA158 used as the GS expression host.

After proteolytic resistant test, the activity of GT and GS was highly decreased and disappeared by all of protease tested, and GA had high protease tolerance as reported that peptides and proteins having cyclic structures are generally stable to treatments of heat, denaturants, and proteases ^30^. These characteristics would be available to application to foods at various stages by combination of GA and GT/GS. Although most cleavage sites of pepsin (FYWML) are overlapping to those of alpha-chymotrypsin (FYW) and proteinase K (FYWMC), the affect of each protease treatment on activity of GA, GT, and GS was different (Table 7, Fig. 3), indicating that the amount of protease units and tertiary structures of gassericins may be important rather than cleavage sites for maintaining the bacteriocin activities. The elucidation of antibacterial spectra, mode of action, and immunity system for GS are in progress.

**Fig. 3.**
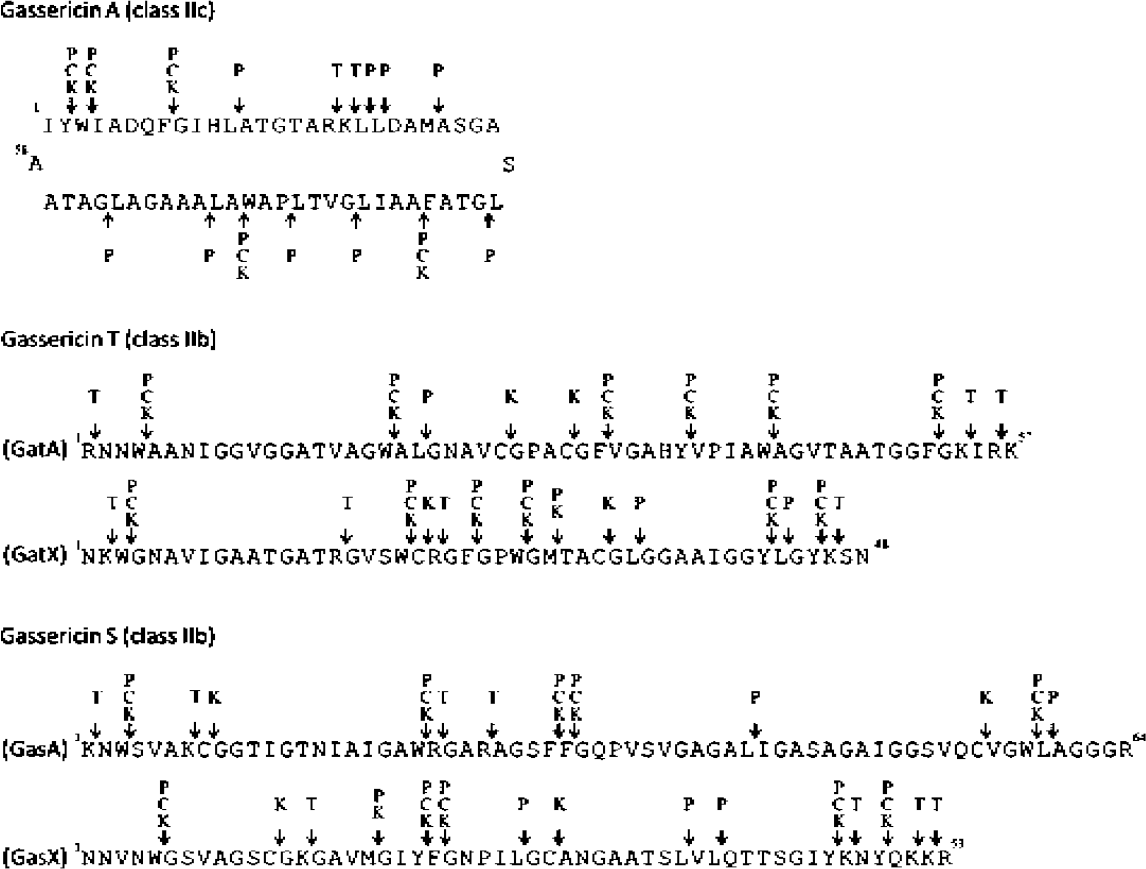
Cleavage sites of each protease on gassericin A, T, and S. Allows indicate the cleavage sites of each protease (P: pepsin, C: alpha - chymotrypsin, K: proteinase K, and T: trypsin)

## Materials and Methods

### Bacterial strains, plasmids, and media

The strains and plasmids used in this study are summarized in Table 8. Gassericin T (GT) producer, *Lactobacillus gasseri* LA158, was isolated from the human feces of a six-month old female infant. Gassericin S (GS) and GT producer, *Lb. gasseri* LA327, was isolated from the human large intestine of an adult. *Lb. gasseri* strains and the indicator strain, *Lb. delbrueckii* subsp. *bulgaricus* (*Lb. bulgaricus*) JCM 1002^T^ (pSYE2), were cultivated in MRS broth (Becton Dickson, MD, USA) and grown at 37 °C. Intermediate expression host strains, *Lactococcus lactis* subsp. *cremoris* (*Lc. cremoris*) MG1363 and *Lactococcus lactis* subsp. *lactis* (*Lc. lactis*) IL1401, were cultivated in M17 medium (Oxoid, Hants, UK) supplemented with 0.5% glucose (GM17) and grown at 30 °C. For plasmid harbours strain, erythromycin (Em) was used as a screening agent in GM-17 broth and MRS broth at final concentration of 25 μg/mL^−1^. All bacteria were stored at −80 °C in their respective media with 30% (w/v) glycerol and cultivated with 10% inoculation more than twice before use.

**Table 8.** Bacterial strains and plasmids used in this study.

| Strain and plasmid | Description<sup>a</sup> | Reference and source<sup>b</sup> |
| --- | --- | --- |
| **Strains** |  |  |
| *Lactobacillus gasseri* |  |  |
| LA39 (JCM 11657) | Producer strain of gassericin A | Laboratory collection |
| LA158 (JCM 11046) | Producer strain of gassericin T, plasmid free | Laboratory collection |
| LA327 | Producer strain of gassericin T and gassericin S, plasmid free | Laboratory collection |
| LA158ΔgatAX | Host strain for recombinant plasmids, plasmid free, derivative of *Lactobacillus gasseri* LA158 lacking the operon *gatAX* | This study |
| LA158ΔgatX | GatA producer strain, plasmid free, derivative of *Lactobacillus gasseri* LA158 lacking the operon *gatX* | This study |
| LA158ΔgatA | GatX producer strain, *Lactobacillus gasseri* LA158 lacking the operon *gatA* | This study |
| LA327ΔgatAX | Producer strain of gassericin S, plasmid free, derivative of *Lactobacillus gasseri* LA327 lacking the operon *gatAX* | This study |
| LA327ΔgatT | Plasmid free, derivative of *Lactobacillus gasseri* LA327 lacking the operon *gatT* | This study |
| *Lactobacillus delbrueckii* subsp. *bulgaricus* JCM1002<sup>T</sup> | Indicator strain for bacteriocin activity assay | JCM |
| *Lactococcus lactis* subsp. *cremoris* MG1363 | Intermediate host strain for recombinant plasmids, plasmid free, derivative of *Lactococcus lactis* subsp. *cremoris* NCDO 712 | Gasson J.M. *et al.* (1983)<sup>40</sup> |
| *Lactococcus lactis* subsp. *lactis* IL1403 | Intermediate host strain for recombinant plasmids, plasmid free | Chopin A. *et al.* (1984)<sup>41</sup> |
| **Plasmids** |  |  |
| pSYE2 | Em<sup>r</sup>; pSY1 derivative with erythromycin resistance gene (*emrA*) of pAMβ1 from *Enterococcus faecalis* | Satoh *et al.* (1997)<sup>42</sup> |
| pIL253-P<sub>32</sub> | Em<sup>r</sup>; pIL253 derivative with P<sub>32</sub> promotor | Kemperman *et al.* (2003)<sup>43</sup> |
| pTERM09 | Em<sup>r</sup>; pSY1 derivative with temperature-sensitive mutation in the *repA* | Ito *et al.* (2009)<sup>32</sup> and Biswas *et al.* (1993)<sup>35</sup> |
| pGS-AX | Em<sup>r</sup>; pIL253-P<sub>32</sub> derivative carrying *gasAX* | This study |
| pGS-AXI | Em<sup>r</sup>; pIL253-P<sub>32</sub> derivative carrying *gasAXI* | This study |
| pGS-AXΔA | Em<sup>r</sup>; pGS-AX derivative eliminated *gasA* | This study |
| pGS-AXΔX | Em<sup>r</sup>; pGS-AX derivative eliminated *gasX* | This study |
| pGS-AXIΔA | Em<sup>r</sup>; pGS-AXI derivative eliminated *gasA* | This study |
| pGS-AXIΔX | Em<sup>r</sup>; pGS-AXI derivative eliminated *gasX* | This study |

<sup>a</sup>Em<sup>r</sup>, Erythromycin resistance.682 <sup>b</sup>JCM, Japan Collection of Microorganisms, Tsukuba, Japan.

### DNA sequencing and genetic analysis

The nucleotide sequences surrounding two bacteriocin structural genes (*gatAX* and *gasAX*) presumed to encoding GT and Acd LF221A were determined with primer walking using the chromosomal DNA of *Lb. gasseri* LA327 prepared by the method of Luchansky *et al.* (1991) 31 as the template. The DNA sequencing was performed using the dideoxy chain termination method with a PRISM 3100 Genetic Analyzer (Applied Biosystems, CA, USA) and a BigDye Terminator v1.1 Cycle Sequencing Kit (Applied Biosystems) according to the protocols of the manufacturers.

Open reading frames (ORFs), promoters, and terminators on the determined DNA sequence were deduced using the GENETYX-MAC software (GENETYX Corporation, Tokyo, Japan) and an online genetic analysis site, SoftBerry (https://blast.ncbi.nlm.nih.gov/Blast.cgi). Predicted amino acid sequences of peptides and proteins encoded by the DNA sequences of the ORFs were subjected to homology search using the BLAST program in the DDBJ database (http://blast.ddbj.nig.ac.jp/top-j.html). Cellular localization and transmembrane segments of each peptide and protein were deduced using two online program TMHMM (http://www.cbs.dtu.dk/services/TMHMM/).

### Electrotransformation of bacteria

Electrotransformation of *Lb. gasseri* strains, *Lc. lactis* IL1403, and *Lc. cremoris* MG1363 was performed as described by Ito *et al.* (2009) ^32^ and Holo *et al.* (1989) ^33^, respectively. Transformants were selected by using 25 μg/mL Em.

### Deletion of the GT structural gene (*gatAX*) and ABC-type transporter gene (*gatT*) from *Lb. gasseri*

The primers used in this study are summarized in Table 9. A cloning vector, pTERM13 32, is derived from pSY1 (GenBank Accession No. E05087) that carries a replication protein gene (*repA*) 100% identical to that of pWVO1 ^34^. The temperature-sensitive (ts) mutation known for pWVO1 *repA* ^35^ was transplanted to *repA* of pTERM13 to obtain a novel ts vector pTERM09. In order to remove *gatAX* and *gatT* from the chromosome of *Lb. gasseri* LA158 and LA327, the double-cross-over (DCO) substitution was used. The 5′-flanking 763 bp and the 3′-flanking 770 bp sequences of *gatAX,* or The 5′-flanking 792 bp and the 3′-flanking 791 bp sequences of *gatT* were amplified and joined by using the splice-overlap extension PCR method with the primers ⊿*gatAX* pr1, ⊿*gatAX* pr2, ⊿*gatAX* pr3, and ⊿*gatAX* pr4 as well as ⊿*gatT* pr1, ⊿*gatT* pr2, ⊿*gatT* pr3, and ⊿*gatT* pr4. Similarly, *gatA* and *gatX* were removed from chromosome of *Lb. gasseri* LA158. After the 5′-flanking 763 bp, and the 3′-flanking 767 bp sequences of *gatA*, or the 5′-flanking 770 bp, and the 3′-flanking 770 bp sequences of *gatX* were amplified with primers ⊿ *gatA* pr1, ⊿*gatA* pr2, ⊿*gatA* pr3, and ⊿*gatA* pr4, as well as ⊿*gatX* pr1, ⊿*gatX* pr2, ⊿ *gatX* pr3, and ⊿*gatX* pr4, they were joined by the aforementioned method. The PCR was done using Phusion DNA polymerase (New England Biolabs, MA, USA). The joined fragment was cloned into the unique *Sma*I-site of pTERM09 to construct pLG⊿*gatAX* and pLG⊿*gatT*. These recombinants were electrotransformed to *Lb. gasseri* LA158 and/or LA327 and transformants were selected at a permissible temperature 32 °C on MRS agar plates containing Em. Integration of the recombinants into the chromosomal *gat* locus of *Lb. gasseri* LA158 and/or LA327 were done by cultivating the transformant at 39 °C and DCO resolution of pLG⊿*gatAX* and pLG⊿ *gatT* from the chromosome were induced by cultivating the integrant at 32 °C in MRS medium without Em. Colony-direct PCR with ⊿*gatAX* pr1 and ⊿*gatAX* pr4 or ⊿*gatT* pr1 and ⊿*gatT* pr4 was used to screen *gatAX*- and *gatT-*deletant clones, from which a 1.5 kb and 1.6 kb fragment was amplified, respectively. Each relevant fragment was sequenced to confirm the correctness of the deletion. The *gatAX*- and *gatT-*deletants thus obtained were designated LA158 ⊿*gatAX,* LA327⊿*gatAX,* and LA327⊿*gatT*.

**Table 9.**
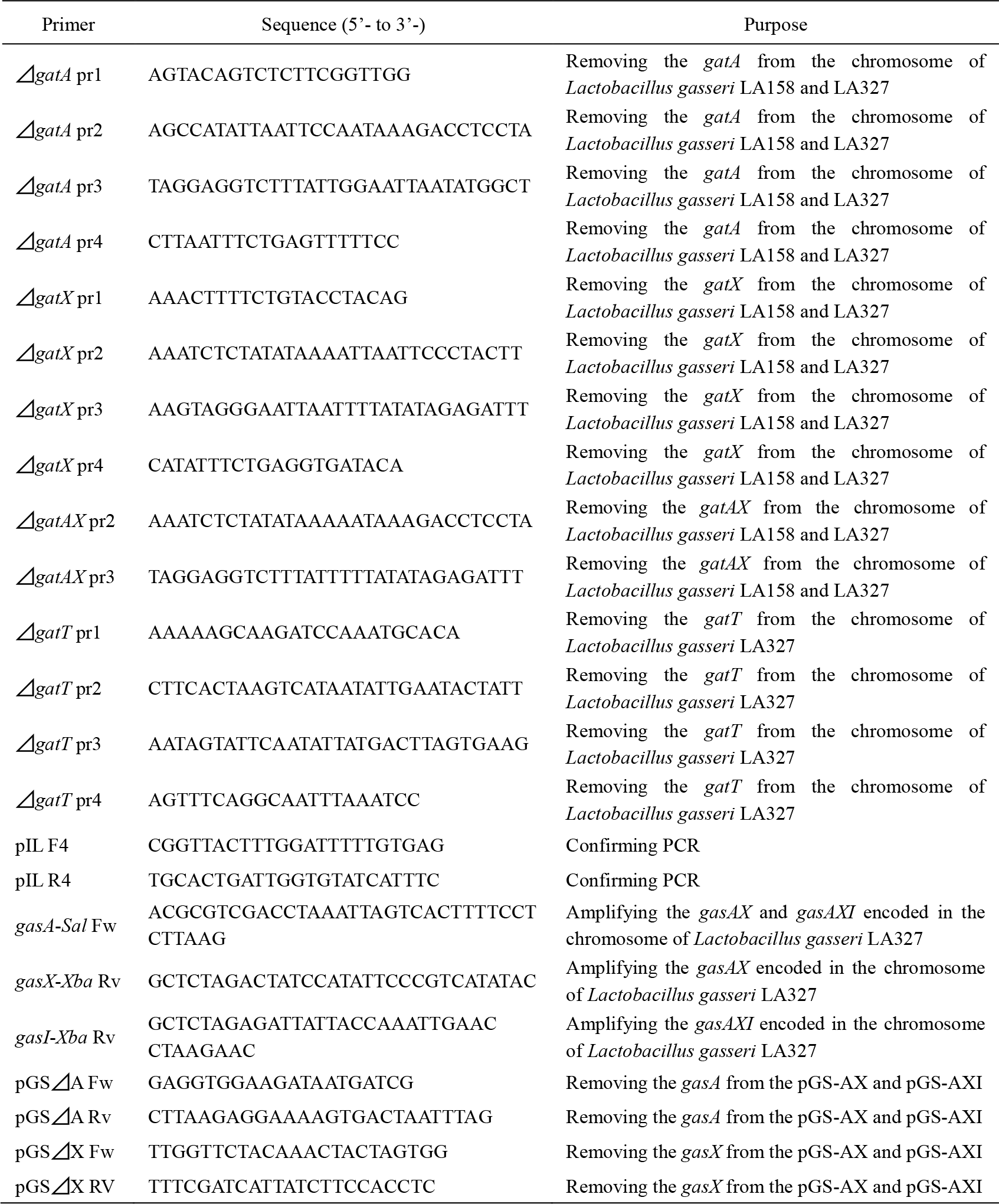
Primers used in this study.

### Construction of GS producing strains for homologous expression

Molecular cloning was performed using standard methods ^36^. The recombinant plasmids of *gasAX* and *gasAXI* (pGS-AX and pGS-AXI, respectively) were constructed in order to compose the GS producers. After *gasAX* and *gasAXI* were amplified by polymerase chain reaction (PCR), using corresponding primers (*gasA-Sal* Fw and *gasX-Xba* Rv for *gasAX*, and *gasA-Sal* Fw and *gasI-Xba* Rv for *gasAXI*), chromosomal DNA from *Lb. gasseri* LA327 as the template DNA, TaKaRa Ex Taq (TaKaRa Bio Inc., Shiga, Japan; EC 2.7.7.7), and T100™ thermal cycler (Bio-Rad Laboratories, Watford, UK). Then, each PCR product was ligated with expression vector (pIL253-P_32_) using T4 DNA Ligase (TaKaRa Bio Inc.; EC 6.5.1.1) via double digestion using *Sal* I and *Xba* I (TaKaRa Bio Inc.; EC 3.1.21.4). Furthermore, to compose the GasA and GasX producers, recombinant plasmids of *gasA, gasX, gasAI,* and *gasXI* (pGS-A, pGS-X, pGS-AI, and pGS-XI, respectively) were constructed by deletion of *gasA* or *gasX* region from pGS-AX and pGS-AXI. Following this, inverse PCR using outward-facing primers to deletion of each region (the primer pair of pGS⊿A Fw and pGS⊿A Rv for deletion of *gasA*, and that of pGS⊿X Fw and pGS⊿X Rv for deletion of *gasX*) was performed. Inverse PCR products were phosphorylated using T4 Polynucleotide Kinase (TaKaRa Bio Inc.; EC 2.7.1.78) and continuously self-ligated using T4 DNA Ligase (TaKaRa Bio Inc.). Each recombinant plasmid was transformed into an intermediate strain *Lc. lactis* subsp. *cremoris* MG1363 and then subsequently into the final expression host strain *Lb. gasseri* LA158⊿*gatAX*.

### Bacteriocin activity assay (agar well diffusion method)

The culture supernatants of each *Lb. gasseri* strain were prepared by centrifugation (8,000×g, 10 min, 4 °C) of the MRS cultures and filtrated through a 0.45 μm membrane filter (ADVANTEC, Tokyo, Japan). Bacteriocin activity was assayed using the agar well diffusion method ^37^. Samples (culture supernatants) were serially diluted to 1/2^n^ using 0.85% PBS. MRS agar plates (90 mm in diameter and 4 mm thick, 15 mL) containing 1.5% (w/v) agar (Oxoid) were overlaid with a soft agar lawn (0.75% agar, 10 mL, 10 μg/mL Em), inoculated with a one-tenth diluted overnight culture (250 μL) of the indicator strain *Lb. delbrueckii* subsp. *bulgaricus* JCM 1002^T^ (pSYE2). Wells (6 mm in diameter) were cut off from the plates, and then 65 μl of the serially diluted samples was added to each well. The plates were incubated for 18 h at 37 °C. The unit of bacteriocin activity (AU: arbitrary unit) was defined as the reciprocal of the highest dilution inhibiting the growth of the indicator strain.

### Bacteriocin activity assay of Gas producers and the synergistic activities of GT and GS

Bacteriocin activities of Gas producers were assayed by the agar well diffusion method as described above. After the bacteriocin activity in the supernatants of GasAX and GasAXI producer were tested, those of GasA, GasX, GasAI, and GasXI producer were examined alone and in combination (GasA or AI and GasX or XI, 1:1). Then, the activity of Gat producers (producing GatA and/or GatX) was similarly assayed. Furthermore, the supernatants of GatA, GatX, GasA, and GasX producer were subjected to the assay in combination (six patterns of two peptides, and the combination of four peptides) to demonstrate the synergistic activity between GT and GS.

### Preparation of crude GS and *In situ* activity assay

The culture supernatant of GS producer, *Lb. gasseri* LA158⊿*gatAX* (pGS-AXI), was concentrated approximately 40 times by ultrafiltration (centrifugal filter units Amicon® Ultra-100K, Merck Millipore, Tullagreen, Ireland), and then the concentrate (crude GS) was subjected to SDS-PAGE following the Laemmli’s method ^38^ with a 4.5% spacer gel and 20% separating gel. Amersham™ ECL™ Rainbow™ Maker-Low Range (GE Healthcare, Tokyo, Japan) with marker range of 3,500-40,000 Da was used as molecular marker. For detection of bacteriocin activity on the gel, *in situ* activity assay was performed as described by Daba *et al* ^39^. Briefly, half cut gel was put in a petri dish after immobilization with 20% isopropanol-10% acetate and being washed with milli-Q water; then, MRS agar containing the indicator strain was stratified and incubated at 37 °C for 24 h. The bacteriocin activity of GS was observed as a clear zone, and the molecular mass was estimated by electrophoretic mobility.

### pH stability assay of GS

Each pH of the culture supernatants from GS producer, *Lb. gasseri* LA158⊿*gatAX* (pGS-AXI), and non bacteriocin producer (control), *Lb. gasseri* LA158⊿*gatAX*, was adjusted to 2, 4, 7, and 10 with 1 N HCl and 1 N NaOH. Following this, the bacteriocin activity of these supernatants was assayed by the agar well diffusion method, using *Lb. delbrueckii* subsp. *bulgaricus* JCM 1002^T^ (pSYE2) as the indicator.

### Heat stability assay of GS

The culture supernatant of GS producer, *Lb. gasseri* LA158⊿*gatAX* (pGS-AXI), was dispensed to 100 μL in 0.2 mL PCR tube (Bio-Rad), simultaneously autoclaved (121 °C, 15 min) or heated (95 °C, 5 min and 70 °C, 1 h). For confirmation of the conservation, each GS solution was incubated at 37 °C for one week and 4 °C for one month, and then offered to the bacteriocin activity assay.

### Proteolytic resistant test of GS

The 0.2% (w/v) Protease solutions were prepared by dissolution of pepsin (Wako pure chemical industries, Osaka, Japan, EC 3.4.23.1) to 0.2 M HCl-KCl buffer (pH 2), and dissolution of trypsin (Wako, USP units/mg; EC 3.4.21.4), alpha-chymotrypsin (MP Biomedicals, Illkirch, France, 40-50 units/mg; EC 3.4.21.1), and proteinase K (Wako, 590 units/mL; EC 3.4.21.64) to 0.2 M sodium phosphate buffer (pH 8). The supernatants of GA producer (*Lb. gasseri* LA39), GT producer (*Lb. gasseri* LA158), and GS producer (*Lb. gasseri* LA158⊿*gatAX* (pGS-AXI)) were mixed with each protease solution in an equal amount, and were incubated at 37 °C, 5 h in a water bath. These treated supernatants were offered to the bacteriocin activity assay.

### Nucleotide sequence accession number

The 6,935 bp and 1,143 bp DNA sequences containing nine genes (*gat*) and three genes (*gas*) were deposited in the DDBJ database under accession numbers of LC389592 and LC389591, respectively.

## Acknowledgments

This research received no specific grant from any funding agency in the public, commercial, or not-for-profit sectors.

## References

1. Zoetendal EG, von Wright A, Vilpponen-Salmela T, Ben-Amor K, Akkermans AD, de Vos WM. 2002. Mucosa-associated bacteria in the human gastrointestinal tract are uniformly distributed along the colon and differ from the community recovered from feces. Appl Environ Microbiol 68(7):3401–3407. PMID:12089021.

2. Reuter G. 2001. The Lactobacillus and Bifidobacterium microflora of the human intestine: Composition and succession. Curr Issues Intest Microbiol 2:43–53. PMID: 11721280.

3. Dall Bello F, Hertel C. 2006. Oral cavity as natural reservoir for intestinal lactobacilli. Syst Appl Microbiol 29:69–76. https://doi.org/10.1016/j.syapm.2005.07.002.

4. Marin ML, Arroyo R, Jimenez E, Gomez A, Fernandez L, Rodriguez JM. 2009. Cold storage of human milk: effect on its bacterial composition. J Pediatr Gastroenterol Nutr 49:343–348. doi: 10.1097/MPG.0b013e31818cf53d.

5. Matsumiya Y, Kato N, Watanabe K, Kato H. 2002. Molecular epidemiological study of vertical transmission of vaginal Lactobacillus species from mothers to newborn infants in Japanese, by arbitrarily primed polymerase chain reaction. J Infect Chemother 8:43–49. https://doi.org/10.1007/s101560200005.

6. Miyazaki K, Kwase M, Kubota A, Yoda K, Harata G, Hosoda M, He F. 2015. Heat-killed Lactobacillus gasseri can enhance immunity in the elderly in a double-blind, placebo-controlled clinical study. Benef Microbes 6:441–449. https://doi.org/10.3920/BM2014.0108.

7. Ghadimi D, Folster-Holst R, de Vrese M, Winkler P, Heller KJ, Schrezenmeir J. 2008. Effect of probiotic bacteria and their genomic DNA on TH1/TH2-cytokine production by peripheral blood mononuclear cells (PBMCs) of healthy and allergic subjects. Immunobiology 213:677–692. https://doi.org/10.1016/j.imbio.2008.02.001.

8. Nishiyama K, Seto Y, Yoshida K, Kakuda T, Takai S, Yamamoto Y, Mukai T. 2014. Lactobacillus gasseri SBT2055 reduces infection by and colonization of Campylobacter jejuni. PLOS One 9(9):e108827. doi: 10.1371/journal.pone.0108827.

9. Kassaa IA, Hober D, Hamze M, Caloone D, Dewilde A, Chihib NE, Drider D. 2015. Vaginal Lactobacillus gasseri CMUL57 can inhibit herpes simplex type 2 but not Coxsackievirus B4E2. Arch Microbiol 197:657–664. doi: 10.1007/s00203-015-1101-8.

10. Motevaseli E, Shirzad M, Akrami SM, Mousavi AS, Mirsalehian A, and Modarressi MH. 2013. Normal and tumour cervical cells respond differently to vaginal lactobacilli, independent of pH and lactate. J Med Microbiol 62:1065–1072. doi: 10.1099/jmm.0.057521-0.

11. Ogawa A, Kobayashi T, Sakai F, Kadooka Y, Kawasaki Y. 2015. Lactobacillus gasseri SBT2055 suppresses fatty acid release through enlargement of fat emulsion size in vitro and promotes fecal fat excretion in healthy Japanese subjects. Lipids Health Dis 14:20–29. doi: 10.1186/s12944-015-0019-0.

12. Klaenhammer TR. 1993. Genetics of bacteriocins produced by lactic acid bacteria. FEMS Microbiol Rev 12:39–85. PMID: 8398217.

13. Cotter PD, Ross RP, Hill C. 2013. Bacteriocins - a viable alternative to antibiotics? Nat Rev Microbiol. 11:95–105. doi: 10.1038/nrmicro2937.

14. Cotter PD, Hill C, Ross RP. 2005. Bacteriocins: developing innate immunity for food. Nat Rev Microbiol 3:777–788. doi: 10.1038/nrmicro1273.

15. Kawai Y, Saito T, Kitazawa H, and Itoh T. 1998. Gassericin A; an Uncommon Cyclic Bacteriocin Produced by Lactobacillus gasseri LA39 Linked at N - and C - Terminal Ends. Biosci Biotechnol Biochem 62:2438–2440. https://doi.org/10.1271/bbb.62.2438.

16. Kawai Y, Saitoh B, Takahashi O, Kitazawa H, Saito T, Nakajima H, Itho T. 2000. Primary amino acid and DNA sequences of gassericin T, a lactacin F-family bacteriocin produced by Lactobacillus gasseri SBT2055. Biosci Biotechnol Biochem 64:2201–2208. https://doi.org/10.1271/bbb.64.2201.

17. Yasuta N, Arakawa K, Kawai Y, Chujo T, Nakamura K, Suzuki H, Ito Y, Nishimura J, Makino Y, Shigenobu S, and Saito T. 2014. Genetic and biochemical evidence for gassericin T production from Lactobacillus gasseri LA158. Milk Science 63:9–17. https://doi.org/10.11465/milk.63.9.

18. Matijašić BB, Rogelj I, Nes IF, Holo H. 1998. Isolation and characterization of two bacteriocins of Lactobacillus acidophilus LF221. Appl Microbiol Biotechnol 49:606–612. https://doi.org/10.1007/s002530051221.

19. Zorič-Peternel M, Čanžek-Majhenič A, Holo H, Nes IF, Salehian Z, Berlec A, Rogelj I. 2010. Wide-inhibitory spectra bacteriocins produced by Lactobacillus gasseri K7. Probiotics Antimicrob Proteins 2(4):233–240. https://doi.org/10.1007/s12602-010-9044-5.

20. Maldonado-Barragán A, Caballero-Guerrero B, Martín V, Ruiz-Barba JL, Rodríguez JM. 2016. Purification and genetic characterization of gassericin E, a novel co-culture inducible bacteriocin from Lactobacillus gasseri EV1461 isolated from the vagina of a healthy woman. BMC Microbiology 16: 37. https://doi.org/10.1186/s12866-016-0663-1.

21. Majhenič AC, Venema K, Allison GE, Matijašić BB, Rogelj I, Klaenhammer TR. 2004. DNA analysis of the genes encoding acidocin LF221 A and acidocin LF221 B, two bacteriocins produced by Lactobacillus gasseri LF221. Appl Microbiol Biotechnol 63(6):705–714. https://doi.org/10.1007/s00253-003-1424-2.

22. Maldonado-Barragán A, Caballero-Guerrero B, Jiménez E, Jiménez-Díaz R, Ruiz-Barba JL, Rodríguez JM. 2009. Enterocin C, a class IIb bacteriocin produced by E. faecalis C901, a strain isolated from human colostrum. Int J Food Microbiol 133:105–112. https://doi.org/10.1016/j.ijfoodmicro.2009.05.008.

23. Mavrić A, Tompa G, Trmčić A, Rogelj I, Matijašić BB. 2014. Bacteriocins of Lactobacillus gasseri K7 – Monitoring of gassericin K7 A and B genes’ expression and isolation of an active component. Process Biochem 49:1251–1259. http://dx.doi.org/10.1016/j.procbio.2014.04.022.

24. Acedo JZ, Towle KM, Lohans CT, Miskolzie M, McKay RT, Doerksen TA, Vederas JC, Martin-Visscher LA. 2017. Identification and three-dimensional structure of carnobacteriocin XY, a class IIb bacteriocin produced by Carnobacteria. FEBS Lett 591:1349–1359. https://doi.org/10.1002/1873-3468.12648.

25. Ra R, Beerthuyzenf MM, de Vos WM, Saris PE, Kuipers OP. 1991. Effects of gene disruptions in the nisin gene cluster of Lactococcus lactis on nisin production and producer immunity. Microbiology 145:1227–1233. doi: 10.1099/13500872-145-5-1227.

26. Anderssen EL, Diep DB, Nes IF, Eijsink VGH, Nissen-Meyer J. 1998. Antagonistic activity of Lactobacillus plantarum C11: two new two-peptide bacteriocins, plantaricin EF and JK, and the induction factor, plantaricin A. Appl Environ Microbiol 64:2269–2272. PMCID: PMC106311.

27. Oppegård C, Fimland G, Thorbæk L, Nissen-Meyer J. 2007. Analysis of the two-peptide bacteriocins lactococcin G and enterocin 1071 by site-directed mutagenesis. Appl Environ Microbiol 73:2931–2938. doi: 10.1128/AEM.02718-06.

28. Oppegård C, Rogne P, Emanuelsen L, Kristiansen PE, Fimland G, Nissen-Meyer J. 2007. The two-peptide class II bacteriocins: structure, production, and mode of action. J Mol Microbiol Biotechnol 13:210–219. https://doi.org/10.1159/000104750.

29. Zendo T, Koga S, Shigeri Y, Nakayama J, Sonomoto K. 2006. Lactococcin Q, a novel two-peptide bacteriocin produced by Lactococcus lactis QU 4. Appl Environ Microbiol 72(5):3383–3389. DOI: 10.1128/AEM.72.5.3383-3389.2006.

30. Craik DJ, Daly NL, Saska I, Trabi M, Rosengren KJ. 2003. Structures of naturally occurring circular proteins from bacteria. J Bacteriol 185(14):4011–4021. doi: 10.1128/JB.185.14.4011-4021.2003.

31. Luchansky JB, Tennant MC, Klaenhammer TR. 1991. Molecular cloning and deoxyribonucleic acid polymorphisms in Lactobacillus acidophilus and Lactobacillus gasseri. J Daily Sci 74:3293–3302. https://doi.org/10.3168/jds.S0022-0302(91)78515-9.

32. Ito Y, Kawai Y, Honme Y, Arakawa K, Sasaki T, and Saito T. 2009. Conjugative plasmid from Lactobacillus gasseri LA39 that carries genes for production of and immunity to the circular bacteriocin gassericin A. Appl Environ Microbiol 75:6340–6351. doi:10.1128/AEM.00195-09.

33. Holo H, and Nes IF. 1989. High-frequency transformation, by electroporation, of Lactococcus lactis subsp. cremoris grown with glycine in osmotically stabilized media. Appl Environ Microbiol 55:3119–3123. PMCID: PMC203233.

34. Leenhouts, KJ, Tolner B, Bron S, Kok J, Venema G, and Seegers JF. 1991. Nucleotide sequence and characterization of the broad-host-range lactococcal plasmid pWVO1. Plasmid 26:55–66. https://doi.org/10.1016/0147-619X(91)90036-V.

35. Biswas I, Gruss A, Ehrlich SD, and Maguin E. 1993. High-efficiency gene inactivation and replacement system for gram-positive bacteria. J Bacteriol 175:3628–3635. PMCID: PMC204764.

36. Sambrook JF, and Russell DW, ed., 2001. Molecular Cloning: A Laboratory Manual, 3rd ed., Vols 1,2 and 3. Cold Spring Harbor Laboratory Press.

37. Toba T, Toshioka E, and Itoh T. 1991. Lacticin, a bacteriocin produced by Lactobacillus delbrueckii subsp. lactis. Lett Appl Microbiol 12:43–45. https://doi.org/10.1111/j.1472-765X.1991.tb00499.x.

38. Leammli UK. 1970. Cleavage of structural proteins during the assembly of the head of bacteriopharge T4. Nature 227:680–685. PMID: 5432063.

39. Daba H, Pandian S, Gosselin JF, Simard RE, Huang J, and Lacroix C. 1991. Detection and activity of a bacteriocin produced by Leuconostoc mesenteroides.: Appl Environ Microbiol 57:3450–3455. PMCID: PMC183995.

40. Gasson JM. 1983. Plasmid complements of Streptococcus lactis NCDO 712 and other lactic streptococci after protoplast-induced curing. J Bacteriol 154:1–9. PMCID: PMC217423.

41. Chopin A, Chopin MC, Moillo-Batt A, and Langella P. 1984. Two plasmid-determined restriction and modification systems in Streptococcus lactis. Plasmid 11:260–263. PMID: 6087394.

42. Satoh E, Ito Y, Sasaki Y, and Sasaki T. 1997. Application of the extracellular α-amylase gene from Streptococcus bovis 148 to construction of a secretion vector for yogurt starter strains. Appl Environ Microbiol 63:4593–4596. PMID: 9361445.

43. Kemperman R, Jonker M, Nauta A, Kuipers OP, and Kok J. 2003. Functional analysis of the gene cluster involved in production of the bacteriocin circularin A by Clostridium beijerinckii ATCC 25752. Appl Environ Microbiol 69:5839–5848. PMCID: PMC201212.

